# Molecular Changes in Prader-Willi Syndrome Neurons Reveals Clues About Increased Autism Susceptibility

**DOI:** 10.1101/2021.08.09.455700

**Authors:** A. Kaitlyn Victor, Martin Donaldson, Daniel Johnson, Winston Miller, Lawrence T. Reiter

## Abstract

**Background:** Prader-Willi syndrome (PWS) is a neurodevelopmental disorder characterized by hormonal dysregulation, obesity, intellectual disability, and behavioral problems. Most PWS cases are caused by paternal interstitial deletions of 15q11.2-q13.1, while a smaller number of cases are caused by chromosome 15 maternal uniparental disomy (PW-UPD). Children with PW-UPD are at higher risk for developing autism spectrum disorder (ASD) than the neurotypical population. In this study, we used expression analysis of PW-UPD neurons to try to identify the molecular cause for increased autism risk.

**Methods:** Dental pulp stem cells (DPSC) from neurotypical control and PWS subjects were differentiated to neurons for mRNA sequencing. Significantly differentially expressed transcripts among all groups were identified. Downstream protein analysis including immunocytochemistry and immunoblots were performed to confirm the transcript level data and pathway enrichment findings.

**Results:** We identified 9 transcripts outside of the PWS critical region (15q11.2-q13.1) that may contribute to core PWS phenotypes. Moreover, we discovered a global reduction in mitochondrial transcripts in the PW-UPD +ASD group. We also found decreased mitochondrial abundance along with mitochondrial aggregates in the cell body and neural projections of +ASD neurons.

**Limitations:** DPSC derived neuronal cultures used here were immature (3 weeks old), while important for studying the development of the disorder, it will be critical to confirm these mitochondrial defects in more mature neurons or postmortem brain tissue. Our PW-UPD -ASD group included only females, but the sample size in downstream image analysis was increased to include males for the analysis of mitochondrial phenotypes. The ASD diagnostic tool we use is the Social Communication Questionnaire (SCQ), which has been used extensively, but is not the gold standard for ASD diagnosis.

**Conclusions:** The 9 transcripts we identified common to all PWS subtypes may reveal PWS specific defects during neurodevelopment. Importantly, we found a global reduction in mitochondrial transcripts in PW-UPD +ASD neurons versus control and other PWS subtypes. We then confirmed mitochondrial defects in neurons from individuals with PWS at the cellular level. Quantification of this phenotype supports our hypothesis that the increased incidence of ASD in PW-UPD subjects may arise from mitochondrial defects in developing neurons.

## INTRODUCTION

Prader-Willi syndrome (PWS) is a multifaceted neurodevelopmental disorder characterized by hypotonia, hyperphagia, and developmental delay [1]. PWS is caused by a loss of expression for one or more paternally expressed genes in the 15q11.2-q13.1 region (the PWS/AS critical region). Most PWS cases are caused by a paternal interstitial deletions in this region, but a smaller percentage of PWS is caused by the inheritance of two copies of maternal chromosome 15, maternal uniparental disomy (UPD) [2]. Due to imprinted expression of critical genes in the 15q11.2-q13.1 region, a set of normally paternally expressed genes are effectively silenced in neurons including *MAGEL2*, *SNORD115/116*, *SNRPN*, and *SNURF* [3]. PWS caused by UPD (PW-UPD) results in a milder phenotype than PWS caused by a paternal deletion (PW-del). However, UPD cases have a higher risk for autism spectrum disorder (ASD) than typically developing individuals [4–8] and later in life can develop cycloid psychosis [9, 10].

The goal of this study is to identify molecular changes that may confer increased autism incidence in PW-UPD subjects through the analysis of expression differences among neurons from PW-UPD, PW-del and control individuals. We used our large collection of dental pulp stem cell (DPSC) lines to generate neurons in culture for these studies [11, 12]. DPSC are neural crest stem cells that reside inside the pulp cavity of naturally shed “baby teeth” [13–15]. They are multipotent stem cells that have been differentiated to various cell types including neurons, osteocytes, glial cells, and adipocytes [16–18]. In fact, several groups have shown that DPSC-derived neuronal cultures exhibit electrophysiological properties of functional neurons [19–22]. Previously, we established DPSC growth parameters [22], efficacy [23], and similarity to other stem cell systems [12]. In an earlier gene expression study of DPSC neurons from duplication 15q11.2-q13.1 (Dup15q) and Angelman syndrome (AS) deletion subjects, we identified distinct expression patterns indicative of each syndrome [24]. For the molecular studies presented here, we differentiated these stem cells into neuronal cultures using our previously published protocol [11]. After differentiation, RNA sequencing (RNAseq) was performed to define transcriptional differences among PWS subtypes and neurotypical controls. These molecular studies revealed new details about shared gene expression changes in PWS and unique expression defects related to mitochondria in PWS-UPD neurons that may contribute to increased autism risk.

## MATERIALS AND METHODS

### Obtaining Teeth for DPSC Cultures

Neurotypical control teeth were obtained through the Department of Pediatric Dentistry and Community Oral Health at the University of Tennessee Health Science Center (UTHSC). Teeth from children with PWS subtypes were collected remotely by the caregivers of these subjects after confirmation of the underlying genetic diagnosis. Subjects provided informed consent for tooth collection along with a Social Communication Questionnaire (SCQ) to assess ASD status. Tooth pulp was cultured from teeth and cell lines frozen during early passages in our DPSC Repository as previously described [11]. The DPSC Repository and subsequent molecular studies were approved by the UTHSC institutional review board prior to conducting research.

### Generation of Dental Pulp Stem Cell Cultures

DPSC used in this study were isolated and cultured according to our previously described protocol and stored in the DPSC Repository [11]. Briefly, after mincing the dental pulp from inside the tooth cavity, 3 mg/mL Dispase II and 4 mg/mL Collagenase I were added to digest the tissue. Cells were then seeded on poly-D-Lysine coated 12-well plates with DMEM/F12 1:1, 10% fetal bovine serum (FBS), 10% newborn calf serum (NCS), and 100 U/mL penicillin and 100 ug/mL streptomycin (Pen/Strep) (Fisher Scientific, Waltham, MA). Confluent cultures (80%) were passaged with TrypLE™ Express and neuronal differentiation performed only on early passage cells (< passage 4).

### Neuronal Differentiation

DPSC lines were seeded at 20,000 cells/cm^2^ on poly-D-lysine coated plates or chamber slides (Ibidi, Planegg, Germany) with DMEM/F12 1:1, 10% fetal bovine serum (FBS), 10% newborn calf serum (NCS), with 100 U/mL penicillin and 100 ug/mL streptomycin (Pen/Strep). At 80% confluence, the neuronal differentiation protocol was followed as previously published in Kiraly *et al.*, 2009 [21] with an extended maturation phase (3 weeks vs 7 days) [11]. Briefly, epigenetic reprogramming was performed by exposing the DPSC to 10 μM 5-azacytidine (Acros Scientific, Geel, Belgium) in DMEM/F12 containing 2.5% FBS and 10 ng/mL bFGF (Fisher Scientific, Waltham, MA) for 48 hours. Neural differentiation was induced by exposing the cells to 250 μM IBMX, 50 μM forskolin, 200 nM TPA, 1mM db-cAMP (Santa Cruz, Dallas, TX), 10 ng/mL bFGF (Invitrogen, Carlsbad, CA), 10 ng/mL NGF (Invitrogen, Carlsbad, CA), 30 ng/mL NT-3 (Peprotech, Rocky Hill, NJ), and 1% insulin-transferrin-sodium selenite premix (ITS) (Fisher Scientific, Waltham, MA) in DMEM/F12 for 3 days. At the end of the neural induction period, neuronal maturation was performed by maintaining the cells in Neurobasal A media (Fisher Scientific, Waltham, MA) with 1mM db-cAMP, 2% B27, 1% N2 supplement, 30 ng/mL NT-3, and 1X Glutamax (Fisher Scientific, Waltham, MA) for 21 days.

### RNA Sequencing of DPSC-Neurons

Once DPSC-neurons were matured for 3 weeks, total RNA was collected using the Zymo Directzol RNA extraction kit (Zymo, Irvine, CA). Extracted RNA was assayed for quality and integrity using the Agilent Bioanalyzer 6000 pico chip (Agilent, Santa Clara, CA). Only RNA with an RNA Integrity Number (RIN) ≥ 9.0 was used for RNAseq studies. Library preparation and RNAseq was performed by Novogene (Sacramento, CA) using the Illumina platform and paired end reads. 20M reads per sample were collected.

### RNAseq Analysis

FASTQ files from Novogene were analyzed for quality and trimmed using FASTQC. All reads were trimmed to remove nucleotides with Phred scores < Q20. The trimmed FASTQ files were aligned to the human genome reference library hg19 using RNASTAR [25]. Once aligned, the SAM files were collected and mined for read count information of each gene present in the reference file. Read counts were normalized using Counts per Million (CPM) method across groups for the entire experiment. Principle component analysis and Pearson’s coefficient plots were performed on the normalized transcriptome profile. A Wilcoxon’s *t*-test was used to determine significance across groups. All genes that fail to yield a p-value ≤0.05 and a fold change greater than 1.5 were removed. Benjamini and Hochberg false discovery rate (FDR) was performed on this trimmed gene list. All genes that failed to yield an FDR rate of ≤ 0.05 were removed. The final significant differential gene lists were loaded into the ClustVis web tool to generate heatmaps [26]. Additionally, the targets were loaded into the web based enrichment analysis tool, Database for Annotation, Visualization and Integrated Discovery (DAVID) [27] to identify enriched gene ontology (GO) terms.

### Immunofluorescence

DPSC were grown and differentiated on 3-well chamber slides (Ibidi, Planegg, Germany) coated with poly-D-lysine. Cells were fixed using a 4% paraformaldehyde solution for 10 minutes. Once fixed, cells were blocked and permeabilized using PBS with 1% BSA, 10% FBS, and 0.3% Triton X-100 for 1 hour. Primary antibodies were diluted to 1:500 for α-Beta Tubulin (Millipore, ab9354) and α-TOMM20 (Santa Cruz, sc-17764) in the blocking solution and incubated overnight at 2-8 °C with agitation. After overnight incubation, the slides were washed 3x with PBS-T for 10 minutes before the secondary antibodies, Goat α-mouse Alexa Fluor 488 (Life Technologies, A11029) and Goat α-chicken Alexa Fluor 594 (Life Technologies, A11042) were added at a 1:1000 dilution. Slides were incubated at room temperature for 1 hour and then washed again 3x for 10 minutes each. Finally, Prolong Gold Antifade with DAPI (Fisher Scientific, Waltham, MA) was applied for mounting. Slides were imaged on a Zeiss 710 confocal microscope at 63x magnification using Z-stacking to image the entire neuron via ZEN software (Black Edition).

### Western Blots

Western blots were performed as previously described [24]. Briefly, protein was extracted from 3-week mature neurons using neuronal protein extraction reagent (N-PER) (Fisher Scientific, Waltham, MA) and protease inhibitor cocktail (Roche). Samples were resolved on a NuPage 1.5mm 4-12% Bis-Tris gel (Invitrogen, Carlsbad, CA) according to manufacturer instructions and transferred to an Invitrolon™-PVDF membrane (Invitrogen, Carlsbad, CA). The membrane was blocked using Odyssey Blocking Buffer (Licor, Lincoln, NE) for 1 hour and incubated overnight at 2-8°C in primary antibody with agitation. Primary antibodies used: α-MAP2 (Santa Cruz, sc-32791), α-Nestin (Santa Cruz, sc-23927), and α-GABA A receptor (Abcam, ab98968). α-GAPDH (Abcam, ab157156) was used as a protein loading control. Blots were incubated at room temperature for 1 hour in secondary antibodies for both the 700 and 800 channels using Li-Cor IR secondary antibodies (Licor, 926-32212 & 926-68074). Blots were imaged on a Li-Cor Odyssey™ Fc Imager. Both the 700 and 800 channels were exposed for 2 minutes.

### Image Analysis

All imaging analysis was done using coded cell lines, so the observer was blinded to the genotypes. Only after data collection and analysis were cell lines decoded. For each cell line (≥5 individuals per group), ≥15 neurons were imaged for analysis of mitochondrial abundance within the neuron. To quantify mitochondrial area, cells were labeled with α-Beta Tubulin (total neuronal area) and α-TOMM20 (mitochondrial area), images were loaded into the Imaris imaging analysis software (Oxford Instruments, Abingdon, UK) and surfaces for each marker were created (see **Fig. 6**). The mitochondrial area (α-TOMM20) was then divided by the total neuronal area (α-Beta Tubulin) to give the percentage of the neuron that contains mitochondria. After collecting all data, significance testing was performed by both ordinary one-way ANOVA and Turkey’s multiple comparisons test with a single pooled variance.

## RESULTS

### PW-UPD subjects have an increased incidence of ASD versus PW-del and control subjects

Several studies have shown an incidence of about 35% for ASD in PW-UPD subjects compared to 18% in PW-del subjects [7, 8, 28]. To measure ASD remotely, we used the Social Communication Questionnaire (SCQ). The SCQ provides a way to ascertain the likelihood a subject may have ASD, especially when face to face evaluation is not possible. Several groups have now validated the SCQ as a screening tool for ASD against other tools like the Autism Diagnostic Interview, Revised (ADI-R) [29–33]. We use the “lifetime” version of the test which has been proven to be a more accurate predictor of ASD [31]. A cutoff score of 15 is considered “possible ASD” according the SCQ lifetime tool. We selected 4 lines from each group (neurotypical control, PW-del, PW-UPD −ASD, and PW-UPD +ASD) for RNAseq analysis and additional lines were used for the immunofluorescence analysis. **Figure 1** shows the SCQ scores from these subjects, except for two neurotypical control subjects for which we did not receive an SCQ. For one of the neurotypical control subjects without SCQ data, we did collect the Social Responsiveness Scale (SRS) which revealed the subject did not have any autism characteristics. The PW-UPD group shows two distinct clusters, those scoring above 15 (possible ASD) and those within normal limits (**Fig. 1**).

**Figure 1.**
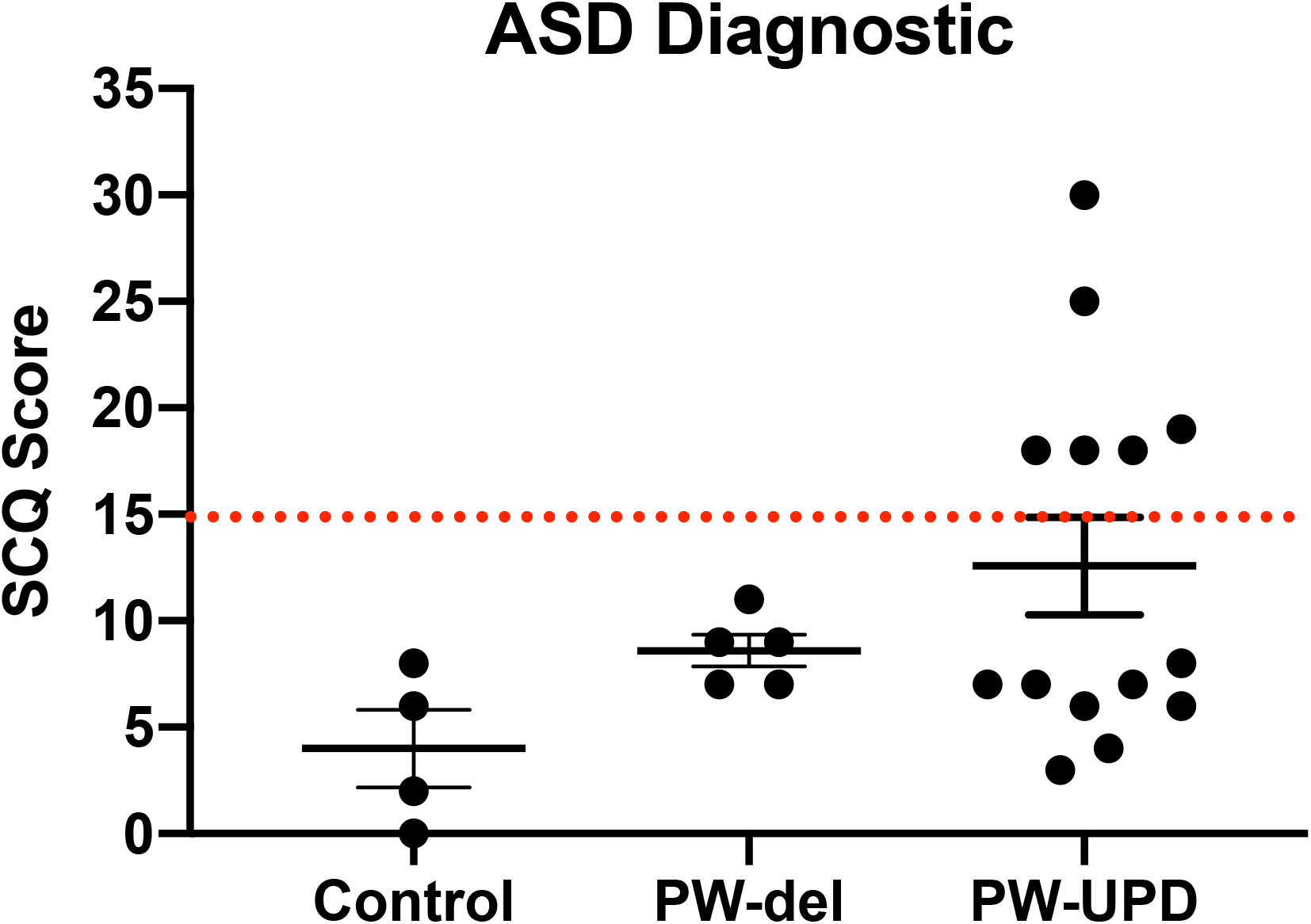
Social Communication Questionnaire (SCQ) scores across all subjects. A cutoff score of 15 (red line) is used as a threshold for “possible ASD”. The PW-UPD cohort has two segregated groups. Subjects that meet the criteria for “Possible ASD” (score of 15 or above) and those scoring within normal limits. For 2 of the neurotypical control subjects used, SCQs could not be collected.

### Characteristics of mixed neuronal cultures derived from control and PWS DPSC lines

Four DPSC lines representing each of the PWS subgroups (PW-del, PW-UPD -ASD, and PW-UPD +ASD) and 4 neurotypical control subject lines were selected from the repository for RNAseq studies (**Supplemental Table 1**). Deciduous (“baby”) teeth from neurotypical control subjects were collected locally from the Department of Pediatric Dentistry and Community Oral Health at UTHSC and PWS subjects were collected remotely by parents and guardians of the subjects using our previously published protocols [11]. DPSC lines were differentiated into neurons using a published neuronal differentiation protocol [11, 21]. DPSC from all groups show positive expression for the stem cell marker, NESTIN, while these same lines were negative for neuronal markers MAP2 and GABA receptor subunit (**Fig. 2A**). After the neuronal differentiation protocol and a 3-week maturation period, the cultures were positive for both MAP2 and GABA A receptor subunit (**Fig. 2A**). DPSC show a flat, fibroblast like morphology (**Fig. 2B**), but after a month of differentiation, the cultures display a pyramidal neuron-like morphology (**Fig. 2C**). We have previously established that a 3-week neuronal maturation period is sufficient to induce significant gene expression changes indicative of terminal neuronal differentiation [22] and reflect underlying disease specific gene expression patterns [24].

**Figure 2.**
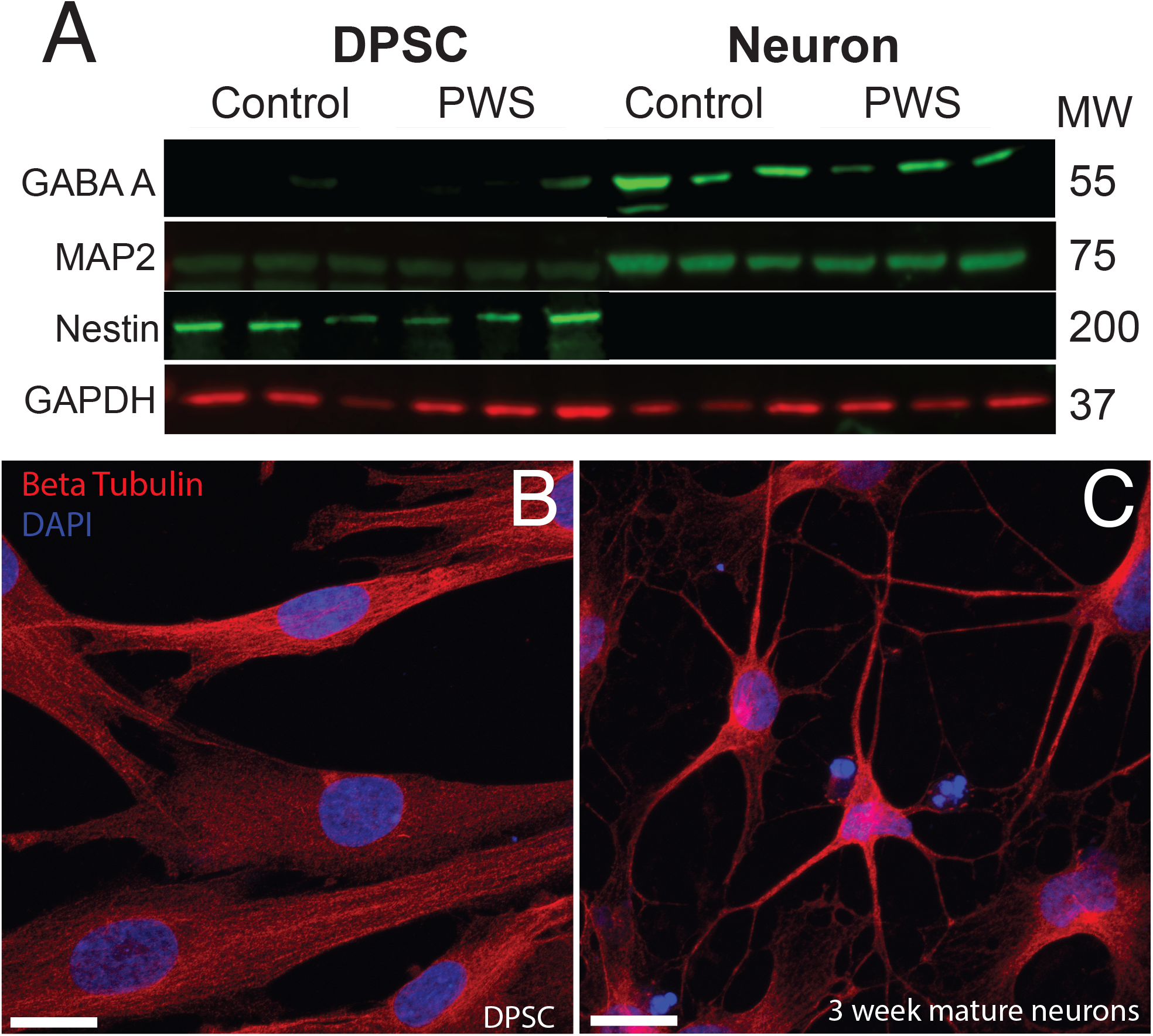
DPSC from both PWS and control subjects differentiate efficiently into neuronal cultures. (**A**) DPSC from all subjects are negative for neuronal markers, MAP2 and GABA A, but positive for stem cell marker NESTIN. After differentiation and 3 weeks of maturation, the cultures are positive for both MAP2 and GABA A receptor subunits. (**B&C**) Representative DPSC vs DPSC-derived neurons visualized with α-beta tubulin (red) (**B**) DPSC show a flat fibroblast-like morphology as previously reported. (**C**) DPSC-derived neuronal culture contains cells with pyramidal neuron morphology. DAPI (blue) was used as a nuclear stain. 63x magnification. Scale bar is 20μM.

### PWS subtypes show distinct transcriptional profiles and a shared PWS expression signature that extends to genes outside of the PWS critical region (15q11.2-q13.1)

RNAseq files from each individual were analyzed by the UTHSC Bioinformatics Core to create lists of significantly (p-value ≤ 0.05 and FDR ≤ 0.05) different transcripts for each PWS subtype versus control and each subtype vs. the other genotypes (see methods for details). Using these lists of significantly differentially expressed transcripts for each group versus control, we created Venn diagrams in BioVenn [34] (**Fig. 3A**). From this analysis we identified a unique molecular signature for each subgroup and a core PWS signature comprised of 3 transcripts within the PWS critical region (*SNRPN*, *SNURF*, and *MAGEL2*) and 9 transcripts coding for genes located outside of 15q PWS/AS critical region (**Fig. 3B**). **Supplemental Table 2** lists these PWS specific transcripts, their function, and evidence that ties these genes or their function to PWS phenotypes. Analyzing the expression of genes across the PWS/AS critical region in our subjects shows the expected expression changes from control to PWS subtypes (**Supplemental Fig. 1**). Specifically, expression of genes that are maternally imprinted in PWS (*MAGEL2*, *SNRPN*, *SNURF*, *NDN*, and *MRKN3*) are absent in our PWS groups, regardless of genetic subtype. These data confirm that DPSC neurons from subjects accurately recapitulate the genetic landscape of PWS in terms of imprinted gene expression.

**Figure 3.**
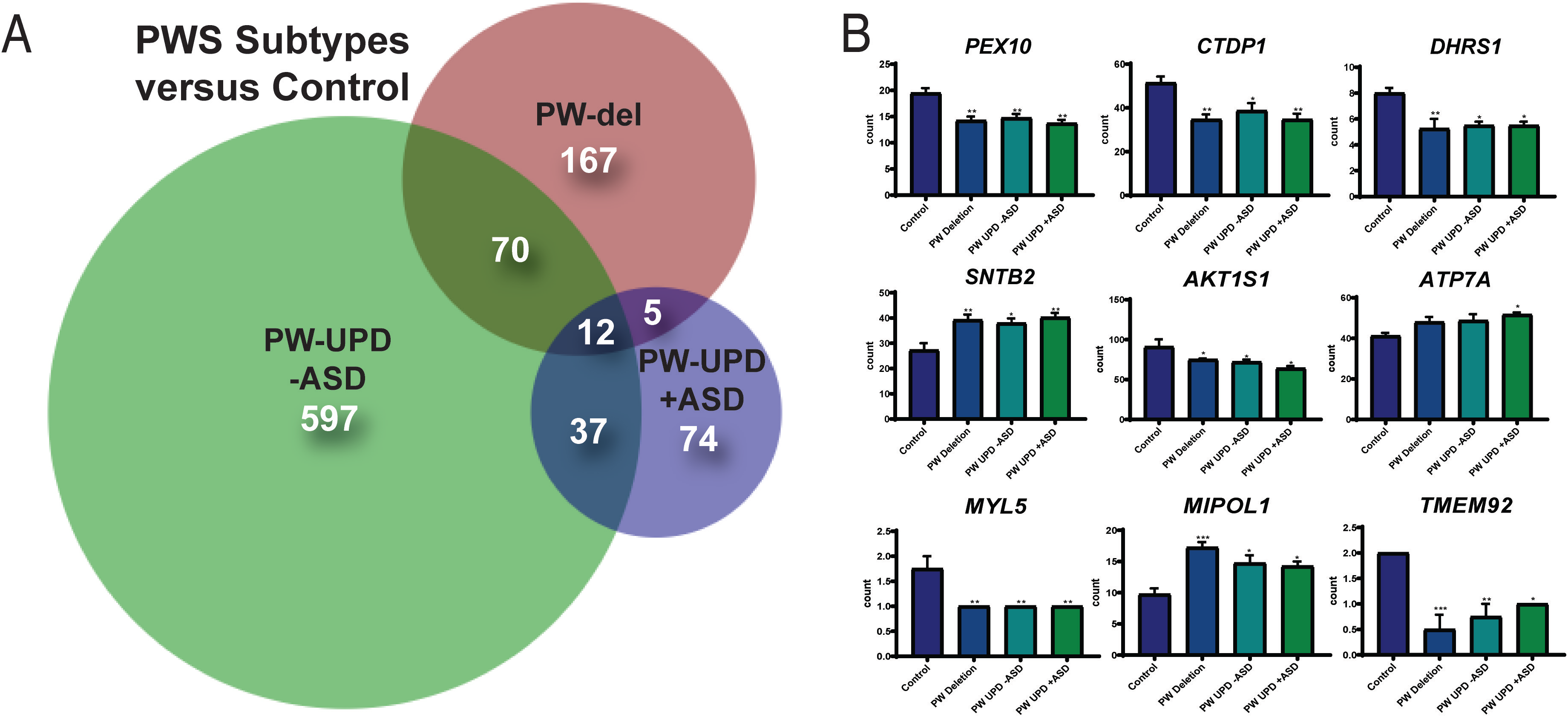
DPSC-derived neuronal cultures define PWS subtype specific expression and a core PWS molecular signature. (**A**) Venn diagram using all the significantly differentially expressed transcripts (p-value ≤ 0.05, FDR ≤ 0.05) versus control for each PWS subtype created using BioVenn [34]. (**B**) Individual bar graphs showing mean counts per transcript across all groups for each of the transcripts located outside of the PWS critical region and identified as common across all PWS subtypes. These 9 genes, along with *MAGEL2, SNRPN,* and *SNURF*, represent a core molecular signature for PWS associated changes in neurons. Significance was determined by individual *t*-tests versus control for each subgroup (p ≤ 0.05).

### Significant Down Regulation of Mitochondrial Transcripts in PWS UPD +ASD Neurons

In order to find molecular changes specific to ASD in our dataset, we investigated the significantly different transcripts (p-value ≤ 0.05, FDR ≤ 0.05) in the −ASD groups versus the UPD +ASD group (**Fig. 4A**). 380 transcripts met our significance cut off and were interrogated for enrichment analysis using the <u>D</u>atabase for <u>A</u>nnotation, <u>V</u>isualization and <u>I</u>ntegrated <u>D</u>iscovery (DAVID), which assigns enrichment scores for Gene Ontology (GO) and other descriptive terms significantly over-represented in our differentially expressed dataset as compared to the entire human genome [27, 35]. Using DAVID functional clustering for GO terms, we found significant enrichments in mitochondrial compartments, functions, and processes (p-value ≤ 0.002) (**Fig. 4B**). The top enrichment clusters were mitochondrial membrane transcripts and transcripts involved in mitochondrial maintenance and biogenesis. It is important to note that no other enrichments (score ≥ 3.0) were found in this dataset. From the transcripts identified in the DAVID enrichment analysis, we created a heatmap across all groups for the mitochondrial genes responsible for the enrichment scores using ClustVis [26] (**Fig. 4C).** This heatmap illustrates that most mitochondrial transcripts identified in our enrichment analysis are dramatically decreased in the PW-UPD +ASD group. **Supplemental Table 3** lists these transcripts and their function. These data support our premise that mitochondrial dysfunction may contribute to increased ASD risk in PW-UPD sub-group.

**Figure 4.**
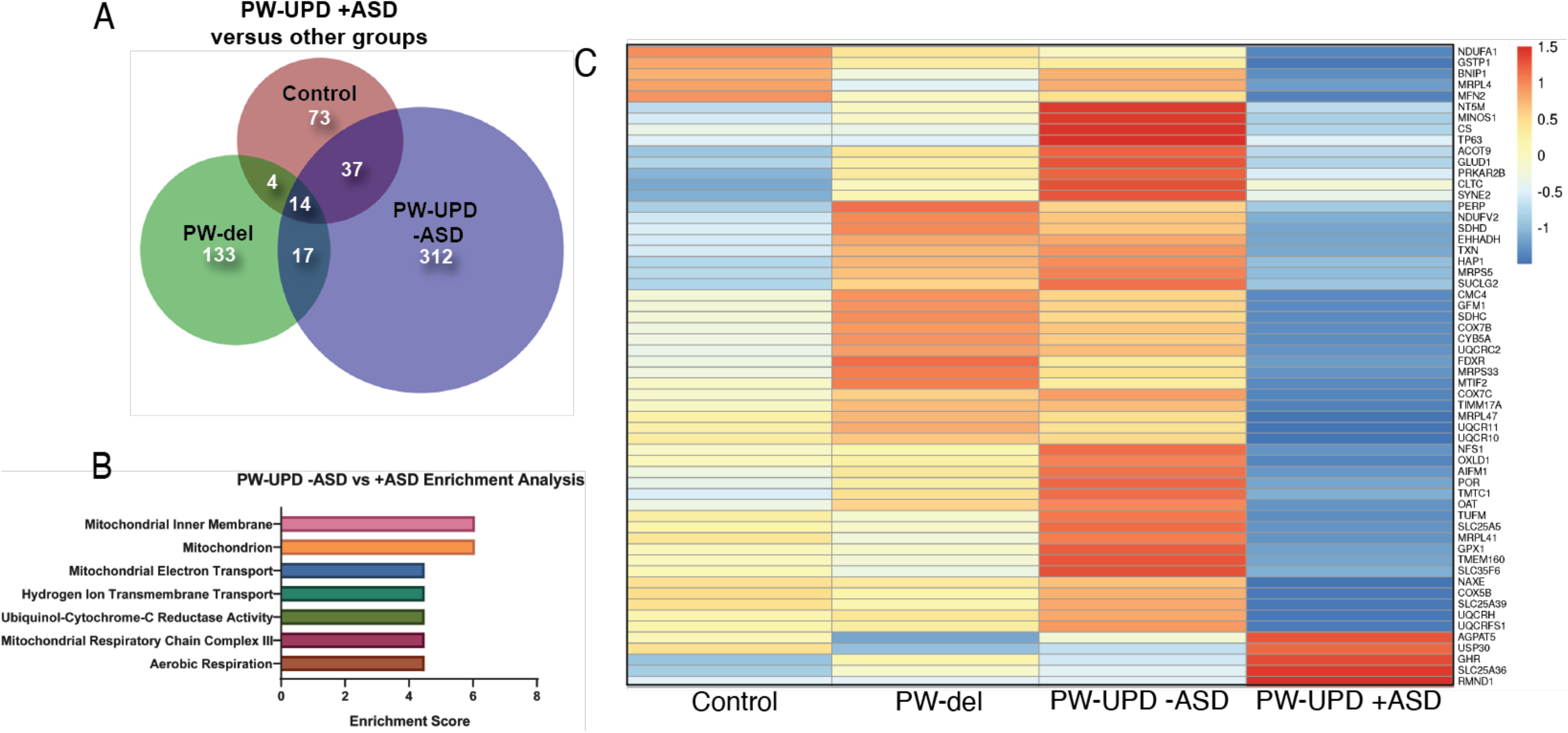
RNAseq analysis reveals enrichment in differentially expressed mitochondrial transcripts in the PW-UPD +ASD group. (**A**) Venn diagram using all the significantly differentially expressed transcripts (p-value ≤ 0.05, FDR ≤ 0.05) versus PW-UPD +ASD for each PWS subtype and control created using BioVenn [34]. (**B**). The list of differentially expressed transcripts PW-UPD −ASD vs UPD +ASD (p-value ≤ 0.05, FDR ≤ 0.05) was used as input for DAVID [27, 35] pathway analysis. The top enrichments (enrichment score ≥ 3.0) were transcripts located within mitochondria and having mitochondrial functions (p-value ≤ 0.002). (**C**) Heatmap of transcripts identified by DAVID analysis as mitochondrial related. Expression of genes identified as driving the mitochondria enrichment in the PW-UPD +ASD group were used to create this heatmap across all subgroups [26]. Colors indicate read counts from high (red) to low (blue). Most of the mitochondrial transcripts identified in DAVID enrichment set have reduced expression in the PW-UPD +ASD group compared to all other groups.

### PW-UPD +ASD Neurons Display Cellular Level Defects in Mitochondria

Since enrichment analysis revealed global down regulation of mitochondrial transcripts, we looked for mitochondrial defects at the cellular level in PW-UPD +ASD neurons. α-Beta Tubulin was used to outline the area of the neurons (**Fig. 5A&C**). To visualize mitochondria in neurons, we used the mitochondrial surface marker α-TOMM20 (translocase of outer mitochondrial membrane 20). α-TOMM20 revealed a mitochondrial phenotype within the +ASD neurons characterized by perinuclear aggregation of mitochondria and decreased mitochondria detected in the neuronal projections (**Fig. 5B vs 5D**). To quantify this mitochondrial specific phenotype, we performed a blinded analysis of neuron images from neurotypical control subjects and PWS subjects. For each group ≥ 5 individuals were used. Each cell line was coded to ensure no user bias when measuring the mitochondrial area within the neuron. The neuronal cultures were immunolabeled with α-TOMM20 and α-Beta Tubulin and then imaged through the cell by confocal microscopy. To determine the percentage of neuronal area containing mitochondria, Imaris software (Oxford Instruments, Abingdon, UK) was used on the confocal stacks. Neuron images were uploaded to the Imaris program and surfaces for quantification were created using the α-Beta Tubulin (**Fig. 6A & 6C**) signal to compute total neuronal area and α-TOMM20 **(Fig. 6E & 6G**) to compute the mitochondrial volume within the neuron. **Figure 6** shows the Imaris output after using the “surfaces” function to define the area of both components. We were able visually to confine our measurements to single neurons and eliminate any debris from this analysis. For each cell line, ≥15 neurons were imaged and analyzed. We found a significant difference in the average neuronal mitochondrial coverage between the PW-UPD +ASD group and the other groups **(Fig. 7)**. These data confirm the hypothesis that mitochondrial dysfunction may underlie the increased ASD incidence of the UPD class.

**Figure 5.**
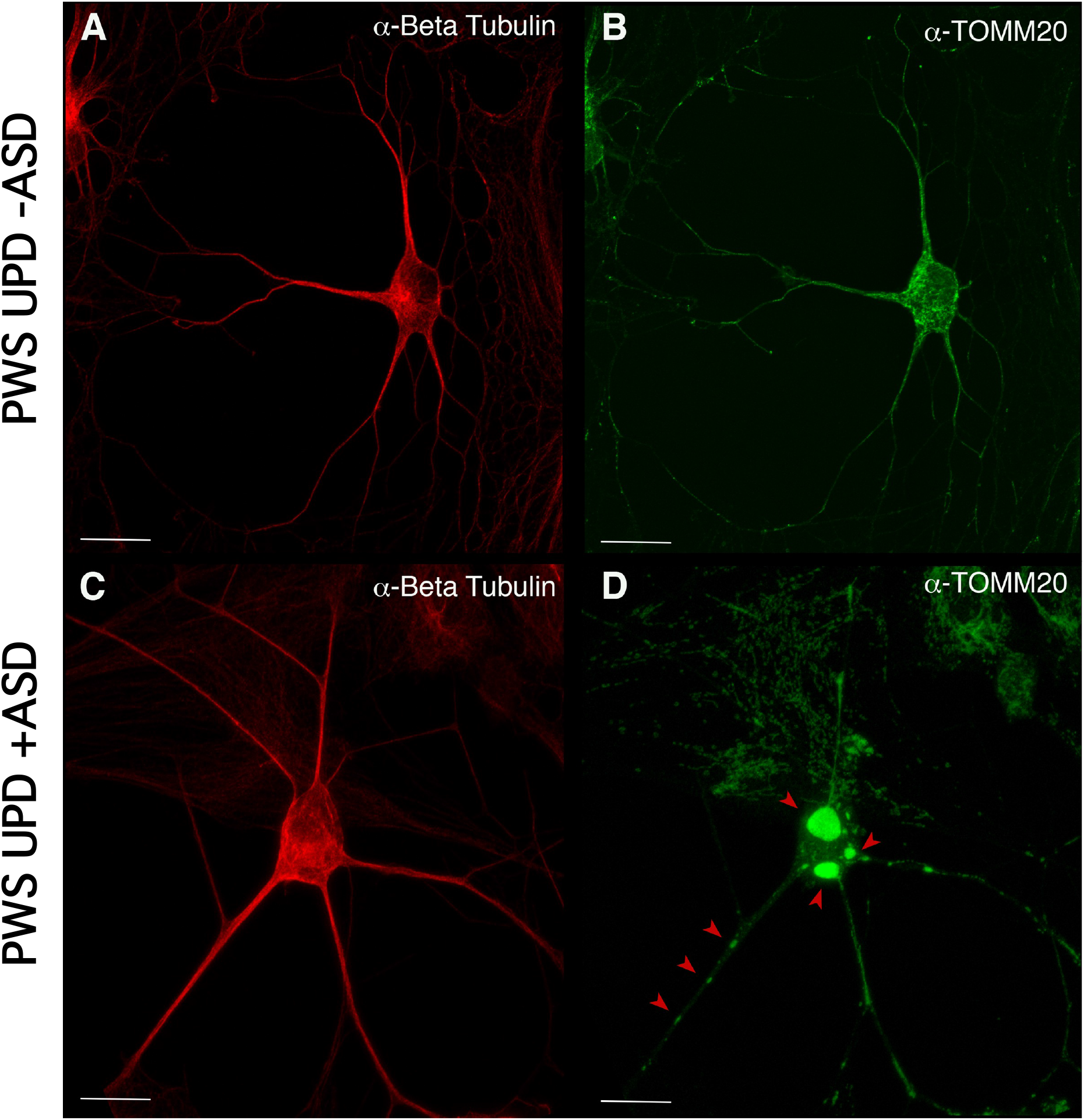
Mislocalization and reduced dispersion of mitochondria in ASD neurons. Neuronal cultures visualized with α-beta tubulin (red) and the mitochondrial marker α-TOMM20 (green). The PW-UPD −ASD neuron (top row) shows bright and evenly dispersed mitochondria within the neuronal projections. In PW-UPD +ASD neuron, the red arrows point to mitochondrial aggregates not seen in the PW-UPD −ASD neurons. **A** & **C** show neuronal morphology using α-beta tubulin staining. **B** & **D** show the α-TOMM20 staining to identify mitochondria. Confocal stacks were taken at 63X magnification. Scale bar is 20μm.

**Figure 6.**
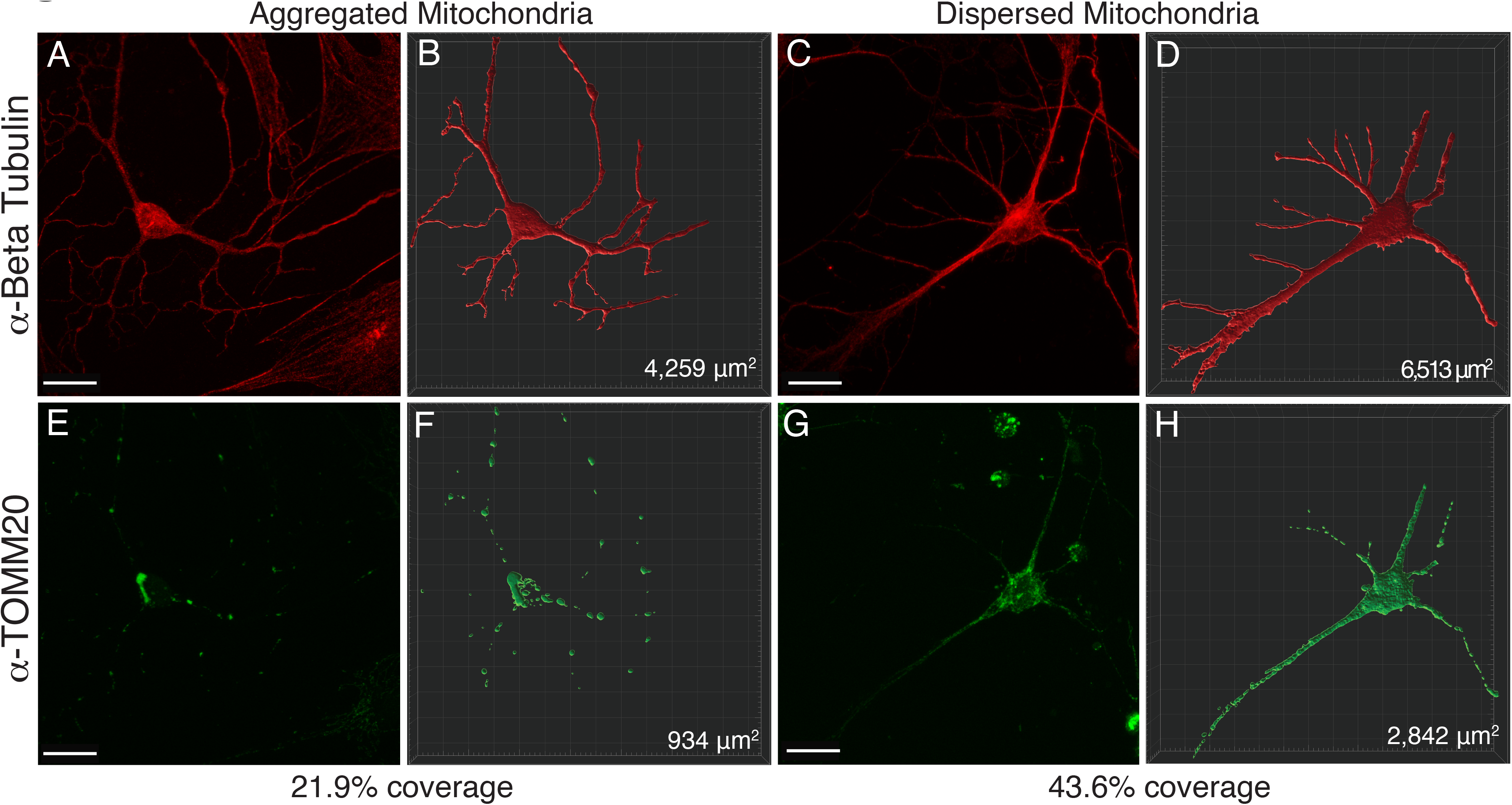
Representative image of Imaris volumetric analysis. Z stacks from neuronal images taken on a Zeiss 810 confocal microscope were loaded into the Imaris software suite for analysis. **A** & **E** show a representative image of a neuron with mitochondrial aggregates while **C** & **G** depict a neuron with dispersed mitochondria. α-Beta Tubulin (**A&C**) was used to measure the surface area of the neuron, while α-TOMM20 (**E&G**) was used to label total mitochondria content within the neuron. Using the surfaces function in Imaris, a mask for the Beta Tubulin (**B&D**) and TOMM20 (**F&H**) stained areas was constructed. From these surface masks, the area is calculated by the software (bottom right corner of **B, D, F** & **H**) for each surface. The mitochondrial area within the neuron was divided by the total neuronal area to determine a percentage of mitochondrial volume. Images were taken at 63x by confocal microscopy. Scale bar represents 20 μM.

**Figure 7.**
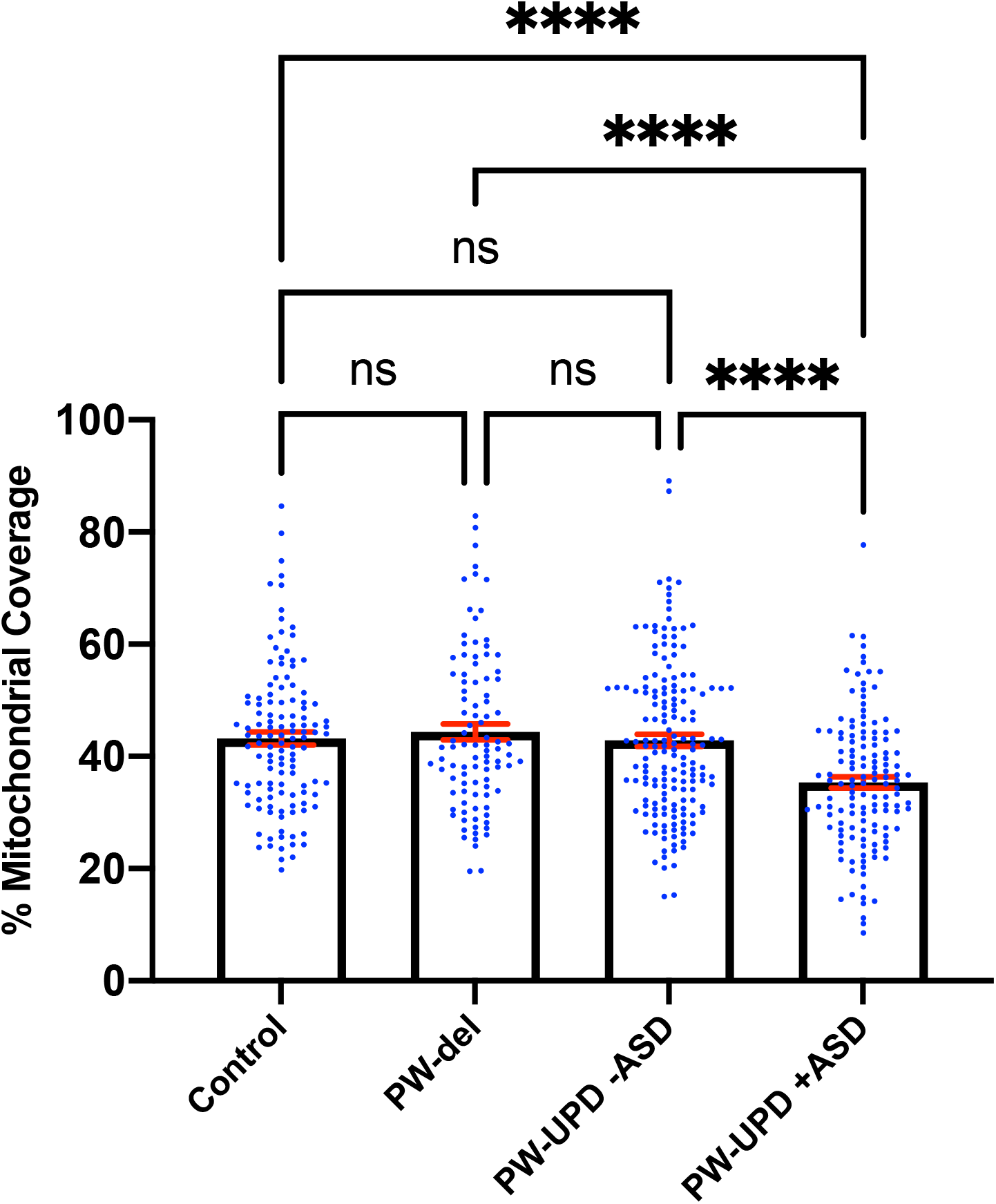
PW-UPD +ASD neurons show decreased mitochondrial coverage within neuronal area. Using Imaris, the total neuronal area (α-Beta Tubulin) that contains mitochondria (α-TOMM20) was calculated. Percent of mitochondrial coverage was measured for ≥15 neurons from each cell line and at least 5 cell lines per group in a blinded fashion. Significance testing was performed by both ordinary one-way ANOVA and Turkey’s multiple comparisons test with a single pooled variance.

## DISCUSSION

Here we used our DPSC-derived neuronal cultures to define molecular signatures for each genomic subtype of PWS, providing for the first time a molecular signature of gene expression in PWS neurons as well as unique molecular changes in the PW-UPD +ASD group that may be indicative of an underlying mitochondrial defect related to autism. DPSC provide an excellent stem cell source to study neurodevelopmental disorders *in vitro* [12]. They are easy to collect remotely and provide a non-invasive way to obtain patient stem cells from a large number of individuals. In addition, DPSC have been found to more closely mimic the epigenetic landscape of embryonic stem cells compared other stem cell types commonly used for neurogenetic research, such as induced pluripotent stem cells (iPSC) [36].

Expression profiling in this study revealed transcripts that may affect early development in PWS, including genes involved in secretory granule regulation (*SNTB2* and *ATP7A*), mTOR signaling (*AKT1S1*), and peroxisome biogenesis (*PEX10*), all of which have been implicated in phenotypes of neurodevelopmental disorders [37–39]. Specifically, prohormone and neuropeptide secretion deficiencies are hallmarks of PWS [40, 41]. These processes rely on efficient secretory granule production and regulation, both of which can be altered by *ATP7A* or *SNTB2,* through its interaction with *PTPRN* [42]. We have previously shown that these MAGEL2 regulated secretory granule defects can be studied in DPSC derived PWS subject neurons. These secretory granule defects have been implicated as a cause of the hormone and neuropeptide deficiencies seen in PWS [41]. *ATP7A* is a copper pump and is integral in regulating copper release at the synapse as well as modulating proteins in the secretory pathway [43]. Due to its widespread function in the brain, it is not surprising that mutations in this gene lead to neurodegenerative and neurodevelopmental disorders [44]. Downstream investigation of these transcripts at the protein level in PWS neurons will be critical to understanding the effect abnormal expression levels of *ATP7A* or *SNTB2* may have on PWS neuropathological development and symptomology. Understanding the function of these genes in the context of PWS could lead to potential therapeutic targets for future interventions involving rescue of endocytic recycling defects driven by loss of MAGEL2 [41].

In addition to uncovering an expression signature common among PWS subtypes compared to neurotypical controls, we also identified transcriptional differences between PW-UPD −ASD and +ASD that may parallel the observed increased autism risk in the PW-UPD cases. In our own cohort of PWS subjects, 33% of PW-UPD individuals scored above the threshold for “possible ASD” on the SCQ (6 out of 18 subjects). This number is comparable to the 35% ASD incidence found in PW-UPD clinical cases [8, 28]. The SCQ has been validated against other ASD tests like ADI-R [30–33, 45] and provides at least an indicator of who may have ASD symptoms.

We identified 380 transcripts that were significantly differentially expressed between +ASD and −ASD groups for PW-UPD subjects (**Fig. 4A**). Enrichment analysis on these 380 transcripts revealed overrepresented GO terms in mitochondrial compartments, functions, and pathways (**Fig. 4B**). Many of the transcripts involved in these enrichments produce proteins that are part of the ETC complexes and proteins involved in mitochondrial function, maintenance and biogenesis (**Supplemental Table 2**) [46, 47]. Cellular analysis in neurons indicated that +ASD neurons showed a mitochondrial aggregation phenotype (**Fig. 5**) and have significantly less mitochondrial volume within the neuron (**Fig. 7**). This further supports our hypothesis that mitochondrial dysfunction (MD) may be involved in the increased ASD incidence in PW-UPD subjects.

MD has been implicated in both idiopathic [48] and syndromic forms of ASD, including Fragile X [49], Down syndrome [50], Rett syndrome [51], and tuberous sclerosis [52]. In fact, it has been reported that up to 80% of individuals with ASD may also have MD [53]. Neurons consume a large amount of energy to maintain ionic gradients and support neurotransmission [54–56] so it is not surprising that mitochondrial defects have been implicated in both neurodevelopmental and neurodegenerative disorders [57–62]. Inside neurons, mitochondria are the main energy producers and are integral in maintaining calcium homeostasis at the synapse, which is vital for regulating neurotransmission [63]. Children diagnosed with both ASD and mitochondrial dysfunction (MD) have a higher rate of neurodevelopmental regression, seizures, and gross motor delay [64, 65]. Several studies have also shown abnormal levels of metabolic biomarkers such as pyruvate, lactate, glutathione, and ubiquinone in ASD subjects [48, 66–68]. Reduced activities in the electron transport chain (ETC), specifically complexes I and V, have been found in the front cortex of ASD individuals [69]. Other studies using post-mortem ASD brain tissue found similar reductions across brain regions of mitochondrial respiratory transcripts and proteins [70–72]. These studies in post-mortem idiopathic ASD brain, although not ideal, add to the evidence that MD may be one of the root causes of ASD in PW-UPD cases.

Performing rescue experiments to re-activate mitochondrial biogenesis, as we did for the restoration of endocytic recycling defects [41], will be a critical future study in our DPSC neuron system. These studies may lead to drug interventions already developed for mitochondrial disorders like Leigh syndrome. In particular, we found a significant decrease in the *PPARGC1A* gene (**Supplemental Fig. 2**) in the PW-UPD +ASD subjects. The protein produced by this transcript, PGC1α, is a transcriptional coactivator that is responsible for regulating mitochondrial biogenesis [73–75]. Using pharmacological agonists already shown to increase PGC1α expression [76, 77], drug screens can be performed to restore the mitochondrial phenotype we found in the PW-UPD +ASD neurons. Coupled with bioenergetic studies of living mitochondria in our PWS neuronal cultures, we will be able to determine the global effects on mitochondrial function caused by paternal loss of expression in PWS subjects.

### Limitations

The neuronal cultures produced from DPSC are immature (3 weeks), which is appropriate for studying neurodevelopmental disorders, but may not reflect disease progression. In addition, the PW-UPD −ASD group consisted of only female subjects, which we recognize could skew our findings. In order to compensate for this, we increased the relative number of male cell lines in the imaging studies, in all groups, to ≥ 5 individuals in order to assess the cellular phenotypes in a more representative number of males and females with PWS. Finally, in order to obtain ASD assessments on our subjects, we relied on the SCQ, which is completed by parents/guardians of PWS subjects. Ideally, we would like to have a more formal ADOS/ADI-R diagnosis [78, 79], but this is not practical for our DPSC collection protocol, and PWS subjects do not often undergo clinical ASD testing as part of their clinical care.

## Conclusions

In this study we utilized our unique collection of patient-derived DPSC lines to generate neurons for next generation RNAseq analysis from three major genomic subtypes of PWS and compared them to neurotypical control neurons. We found 9 genes outside the PWS/AS critical region on 15q11.2-q13.1 that may contribute to the overall PWS phenotype. We also identified an expression signature in the PW-UPD +ASD group compared to all other groups that appears to indicate a global down regulation of mitochondrial genes is associated with ASD in the PW-UPD subjects. Finally, we showed that this molecular signature translated to a visible and quantifiable change in the appearance and abundance of mitochondria in PW-UPD +ASD neurons. These studies are the first steps at investigating the molecular defects in PWS neurons contributing to both PWS symptomology and increased autism incidence in PW-UPD cases.

## List of abbreviations

ASD: Autism spectrum disorder
DAVID: Database for Annotation, Visualization and Integrated Discovery
DPSC: dental pulp stem cells
ETC: electron transport chain
FBS: fetal bovine serum
FDR: false discovery rate
GO: gene ontology
MD: mitochondrial dysfunction
NCS: newborn calf serum
PWS: Prader-Willi syndrome
PW-UPD: PWS by UPD
PW-del: PWS by deletion
SCQ: Social Communication Questionnaire
TOMM20: translocase of the outer mitochondrial membrane 20

## DECLARATIONS

### Ethics approval and consent to participate

The UTHSC Institutional Review Board (IRB) approved this study and informed consent was obtained for both participation and publication of results in accordance with IRB policy.

### Availability of data and materials

The datasets generated and analyzed during the current study are available in the GEO repository (GSE178687). https://www.ncbi.nlm.nih.gov/geo/query/acc.cgi?acc=GSE178687

### Author Contributions

LTR conceived of and acquired funding for this study. AKV grew cultures and performed experiments. DJ and WM performed bioinformatic analysis. AKV and LTR analyzed data and wrote the manuscript.

## Acknowledgments

The authors would like to thank the PWS families who donated teeth to this study. We also thank the Foundation for Prader-Willi Research for advertising this study to potential participants and for their continued support of our work. We also thank the UTHSC Neuroscience Institute and former director of the imaging core, Dr. TJ Hollingsworth for his help with Imaris image analysis.

---

**Supplemental Figure 1. Genes in the 15q11.2-q13 critical region shows expected PWS imprinted expression.** Across the PWS/AS critical region, maternally imprinted genes such as *MAGEL2*, *SNRPN*, and *SNURF* showed decreased expression in both PW-del and PW-UPD neurons.

**Supplemental Figure 2. Expression of mitochondrial biogenesis factor, PPARGC1A, is significantly decreased in PW-UPD +ASD neurons.** Mean RNAseq count for each group is shown in the graph. Significance was determined by an unpaired *t*-test (p ≤ 0.05).

## REFERENCES CITED

1. Cassidy SB, Schwartz S, Miller JL, Driscoll DJ: Prader-Willi syndrome. Genetics in medicine : official journal of the American College of Medical Genetics 2012, 14(1):10–26.

2. Cheon CK: Genetics of Prader-Willi syndrome and Prader-Will-Like syndrome. Ann Pediatr Endocrinol Metab 2016, 21(3):126–135.

3. Angulo MA, Butler MG, Cataletto ME: Prader-Willi syndrome: a review of clinical, genetic, and endocrine findings. Journal of endocrinological investigation 2015, 38(12):1249–1263.

4. Bennett JA, Germani T, Haqq AM, Zwaigenbaum L: Autism spectrum disorder in Prader-Willi syndrome: A systematic review. American Journal of Medical Genetics 2015(167A):2936–2944.

5. Dykens EM, Roof E, Hunt-Hawkins H, Dankner N, Lee EB, Shivers CM, Daniell C, Kim SJ: Diagnoses and characteristics of autism spectrum disorders in children with Prader-Willi syndrome. J Neurodev Disord 2017, 9:18.

6. Whittington J, Holland A: Neurobehavioral phenotype in Prader-Willi syndrome. American journal of medical genetics Part C, Seminars in medical genetics 2010, 154c(4):438–447.

7. Baker EK, Godler DE, Bui M, Hickerton C, Rogers C, Field M, Amor DJ, Bretherton L: Exploring autism symptoms in an Australian cohort of patients with Prader-Willi and Angelman syndromes. Journal of neurodevelopmental disorders 2018, 10(1):24–24.

8. Dimitropoulos A, Zyga O, Russ SW: Early Social Cognitive Ability in Preschoolers with Prader-Willi Syndrome and Autism Spectrum Disorder. J Autism Dev Disord 2019, 49(11):4441–4454.

9. Verhoeven WM, Tuinier S, Curfs LM: Prader-Willi syndrome: cycloid psychosis in a genetic subtype? Acta Neuropsychiatr 2003, 15(1):32–37.

10. Singh D, Sasson A, Rusciano V, Wakimoto Y, Pinkhasov A, Angulo M: Cycloid Psychosis Comorbid with Prader-Willi Syndrome: A Case Series. American journal of medical genetics Part A 2019, 179(7):1241–1245.

11. Goorha S, Reiter LT: Culturing and Neuronal Differentiation of Human Dental Pulp Stem Cells. Curr Protoc Hum Genet 2017, 92(21):21 26 21–21 26 10.

12. Victor AK, Reiter LT: Dental pulp stem cells for the study of neurogenetic disorders. Hum Mol Genet 2017, 26(R2):R166–R171.

13. Gronthos S, Brahim J, Li W, Fisher LW, Cherman N, Boyde A, DenBesten P, Robey PG, Shi S: Stem Cell Properties of Human Dental Pulp Stem Cells. J Dent Res 2002, 81(8):531–535.

14. Gronthos S, Mankani M, Brahim J, Gehron Robey P, Shi S: Postnatal human dental pulp stem cells (DPSCs) in vitro and in vivo. PNAS 2000, 97(25):13625 – 13630.

15. Miura M, Gronthos S, Zhao M, Lu B, Fisher LW, Robey PG, Shi S: SHED: Stem cells from human exfoliated deciduous teeth. PNAS 2003, 100(10):5807–5812.

16. Ikbale E-A, Goorha S, Reiter LT, Miranda-Carboni GA: Effects of hTERT immortalization on osteogenic and adipogenic differentiation of dental pulp stem cells. Data Brief 2016, 6:696–699.

17. Nuti N, Corallo C, Chan BMF, Ferrari M, Gerami-Naini B: Multipotent Differentiation of Human Dental Pulp Stem Cells: a Literature Review. Stem Cell Reviews and Reports 2016, 12(5):511–523.

18. Young FI, Telezhkin V, Youde SJ, Langley MS, Stack M, Kemp PJ, Waddington RJ, Sloan AJ, Song B: Clonal Heterogeneity in the Neuronal and Glial Differentiation of Dental Pulp Stem/Progenitor Cells. Stem Cells International 2016, 2016.

19. Li D, Zou XY, El-Ayachi I, Romero LO, Yu Z, Iglesias-Linares A, Cordero-Morales JF, Huang GT: Human Dental Pulp Stem Cells and Gingival Mesenchymal Stem Cells Display Action Potential Capacity In Vitro after Neuronogenic Differentiation. Stem Cell Rev 2019, 15(1):67–81.

20. Gervois P, Struys T, Hilkens P, Bronckaers A, Ratajczak J, Politis C, Brone B, Lambrichts I, Martens W: Neurogenic maturation of human dental pulp stem cells following neurosphere generation induces morphological and electrophysiological characteristics of functional neurons. Stem Cells and Development 2015, 24(2015):296–311.

21. Kiraly M, Porcsalmy B, Pataki A, Kadar K, Jelitai M, Molnar B, Hermann P, Gera I, Grimm W-D, Ganss B et al: Simultaneous PKC and cAMP activation induces differentiation of human dental pulp stem cells into functionally active neurons. Neurochemistry International 2009, 55:323–332.

22. Urraca N, Memon R, El-Iyachi I, Goorha S, Valdez C, Tran QT, Scroggs R, Miranda-Carboni GA, Donaldson M, Bridges D et al: Characterization of neurons from immortalized dental pulp stem cells for the study of neurogenetic disorders. Stem Cell Research 2015, 15:722–730.

23. Wilson R, Urraca N, Skobowiat C, Hope KA, Miravalle L, Chamberlin R, Donaldson M, Seagroves TN, Reiter LT: Assesment of the tumorigenic potential of spontaneously immortalized and hTERT immortalized cultured dental pulp stem cells. Stem Cells Translational Medicine 2015(4):905–912.

24. Urraca N, Hope K, Victor AK, Belgard TG, Memon R, Goorha S, Valdez C, Tran QT, Sanchez S, Ramirez J et al: Significant transcriptional changes in 15q duplication by not Angelman syndrome deletion stem cell-derived neurons. Molecular Autism 2018, 6(9).

25. Widmann J, Stombaugh J, McDonald D, Chocholousova J, Gardner P, Iyer MK, Liu Z, Lozupone CA, Quinn J, Smit S et al: RNASTAR: an RNA STructural Alignment Repository that provides insight into the evolution of natural and artificial RNAs. Rna 2012, 18(7):1319–1327.

26. Metsalu T, Vilo J: ClustVis: a web tool for visualizing clustering of multivariate data using Principal Component Analysis and heatmap. Nucleic Acids Research 2015, 43(W1):W566–W570.

27. Huang da W, Sherman BT, Lempicki RA: Systematic and integrative analysis of large gene lists using DAVID bioinformatics resources. Nat Protoc 2009, 4(1):44–57.

28. Bennett JA, Germani T, Haqq AM, Zwaigenbaum L: Autism spectrum disorder in Prader-Willi syndrome: A systematic review. American journal of medical genetics Part A 2015, 167a(12):2936–2944.

29. Mouti A, Dryer R, Kohn M: Differentiating Autism Spectrum Disorder From ADHD Using the Social Communication Questionnaire. Journal of attention disorders 2019, 23(8):828–837.

30. Hutchins P, Williams K, Silove N, Allen C: Validity of the social communication questionnaire in assessing risk of autism in preschool children with developmental problems. J Autism Dev Disord 2007, 37(7):1272–1278.

31. Chesnut SR, Wei T, Barnard-Brak L, Richman DM: A meta-analysis of the social communication questionnaire: Screening for autism spectrum disorder. Autism : the international journal of research and practice 2017, 21(8):920–928.

32. Barnard-Brak L, Richman DM, Chesnut SR, Little TD: Social Communication Questionnaire scoring procedures for autism spectrum disorder and the prevalence of potential social communication disorder in ASD. School psychology quarterly : the official journal of the Division of School Psychology, American Psychological Association 2016, 31(4):522–533.

33. Sappok T, Diefenbacher A, Gaul I, Bolte S: Validity of the social communication questionnaire in adults with intellectual disabilities and suspected autism spectrum disorder. American journal on intellectual and developmental disabilities 2015, 120(3):203–214.

34. Hulsen T, de Vlieg J, Alkema W: BioVenn – a web application for the comparison and visualization of biological lists using area-proportional Venn diagrams. BMC Genomics 2008, 9(1):488.

35. Huang da W, Sherman BT, Lempicki RA: Bioinformatics enrichment tools: paths toward the comprehensive functional analysis of large gene lists. Nucleic Acids Res 2009, 37(1):1–13.

36. Dunaway K, Goorha S, Matelski L, Urraca N, Lein P, Korf I, Reiter L, LaSalle J: Dental pulp stem cells model early life and imprinted DNA methylation patterns. Stem Cells 2017.

37. Barone R, Rizzo R, Tabbì G, Malaguarnera M, Frye RE, Bastin J: Nuclear Peroxisome Proliferator-Activated Receptors (PPARs) as Therapeutic Targets of Resveratrol for Autism Spectrum Disorder. Int J Mol Sci 2019, 20(8).

38. Wang L, Zhou K, Fu Z, Yu D, Huang H, Zang X, Mo X: Brain Development and Akt Signaling: the Crossroads of Signaling Pathway and Neurodevelopmental Diseases. Journal of molecular neuroscience : MN 2017, 61(3):379–384.

39. Niciu MJ, Ma XM, El Meskini R, Ronnett GV, Mains RE, Eipper BA: Developmental changes in the expression of ATP7A during a critical period in postnatal neurodevelopment. Neuroscience 2006, 139(3):947–964.

40. Burnett LC, LeDuc CA, Sulsona CR, Paull D, Rausch R, Eddiry S, Carli JF, Morabito MV, Skowronski AA, Hubner G et al: Deficiency in prohormone convertase PC1 impairs prohormone processing in Prader-Willi syndrome. J Clin Invest 2017, 127(1):293–305.

41. Chen H, Victor AK, Klein J, Tacer KF, Tai DJC, de Esch C, Nuttle A, Temirov J, Burnett LC, Rosenbaum M et al: Loss of MAGEL2 in Prader-Willi syndrome leads to decreased secretory granule and neuropeptide production. JCI Insight 2020, 5(17).

42. Suckale J, Solimena M: The insulin secretory granule as a signaling hub. Trends in endocrinology and metabolism: TEM 2010, 21(10):599–609.

43. D’Ambrosi N, Rossi L: Copper at synapse: Release, binding and modulation of neurotransmission. Neurochem Int 2015, 90:36–45.

44. Kaler SG: ATP7A-Related Copper Transport Disorders. In: GeneReviews. Edited by Adam MP, Ardinger HH, Pagon RA, Wallace SE, Bean LJH, Stephens K, Amemiya A. Seattle (WA): University of Washington, Seattle; 1993.

45. Schanding GTJ, Nowell KP, Goin-Kochel RP: Utility of the Social Communication Questionnaire-Current and Social Responsiveness Scale as Teacher-Report Screening Tools for Autism Spectrum Disorders. J Autism Dev Disord 2012, 42.

46. Guo R, Gu J, Zong S, Wu M, Yang M: Structure and mechanism of mitochondrial electron transport chain. Biomed J 2018, 41(1):9–20.

47. Cogliati S, Lorenzi I, Rigoni G, Caicci F, Soriano ME: Regulation of Mitochondrial Electron Transport Chain Assembly. J Mol Biol 2018, 430(24):4849–4873.

48. Weissman JR, Kelley RI, Bauman ML, Cohen BH, Murray KF, Mitchell RL, Kern RL, Natowicz MR: Mitochondrial disease in autism spectrum disorder patients: a cohort analysis. PLoS One 2008, 3(11):e3815.

49. Abbeduto L, Thurman AJ, McDuffie A, Klusek J, Feigles RT, Ted Brown W, Harvey DJ, Adayev T, LaFauci G, Dobkins C et al: ASD Comorbidity in Fragile X Syndrome: Symptom Profile and Predictors of Symptom Severity in Adolescent and Young Adult Males. Journal of Autism and Developmental Disorders 2019, 49(3):960–977.

50. Izzo A, Mollo N, Nitti M, Paladino S, Calì G, Genesio R, Bonfiglio F, Cicatiello R, Barbato M, Sarnataro V et al: Mitochondrial dysfunction in down syndrome: molecular mechanisms and therapeutic targets. Molecular medicine (Cambridge, Mass) 2018, 24(1):2.

51. Hirofuji S, Hirofuji Y, Kato H, Masuda K, Yamaza H, Sato H, Takayama F, Torio M, Sakai Y, Ohga S et al: Mitochondrial dysfunction in dopaminergic neurons differentiated from exfoliated deciduous tooth-derived pulp stem cells of a child with Rett syndrome. Biochemical and biophysical research communications 2018, 498(4):898–904.

52. Ebrahimi-Fakhari D, Saffari A, Wahlster L, Di Nardo A, Turner D, Lewis TL, Jr., Conrad C, Rothberg JM, Lipton JO, Kolker S et al: Impaired Mitochondrial Dynamics and Mitophagy in Neuronal Models of Tuberous Sclerosis Complex. Cell Rep 2016, 17(4):1053–1070.

53. Rose S, Niyazov DM, Rossignol DA, Goldenthal M, Kahler SG, Frye RE: Clinical and molecular characteristics of mitochondrial dysfunction in Autism Spectrum disorder. Molecular Diagnosis & Therapy 2019, 2018(22):517–593.

54. Kann O, Kovacs R: Mitochondria and neuronal activity. Am J Physiol Cell Physiol 2007, 292(2):C641–657.

55. Li Z, Okamoto K, Hayashi Y, Sheng M: The importance of dendritic mitochondria in the morphogenesis and plasticity of spines and synapses. Cell 2004, 119(6):873–887.

56. Princz A, Kounakis K, Tavernarakis N: Mitochondrial contributions to neuronal development and function. Biological chemistry 2018, 399(7):723–739.

57. Rose S, Niyazov DM, Rossignol DA, Goldenthal M, Kahler SG, Frye RE: Clinical and Molecular Characteristics of Mitochondrial Dysfunction in Autism Spectrum Disorder. Mol Diagn Ther 2018, 22(5):571–593.

58. Rojas-Charry L, Nardi L, Methner A, Schmeisser MJ: Abnormalities of synaptic mitochondria in autism spectrum disorder and related neurodevelopmental disorders. Journal of molecular medicine (Berlin, Germany) 2021, 99(2):161–178.

59. Burté F, Carelli V, Chinnery PF, Yu-Wai-Man P: Disturbed mitochondrial dynamics and neurodegenerative disorders. Nat Rev Neurol 2015, 11(1):11–24.

60. Lin MT, Beal MF: Mitochondrial dysfunction and oxidative stress in neurodegenerative diseases. Nature 2006, 443(7113):787–795.

61. Johnson J, Mercado-Ayon E, Mercado-Ayon Y, Dong YN, Halawani S, Ngaba L, Lynch DR: Mitochondrial dysfunction in the development and progression of neurodegenerative diseases. Arch Biochem Biophys 2021, 702:108698.

62. Rossignol D, Frye R: Mitochondrial dysfunction in autism spectrum disorders: a systematic review and meta-analysis. Molecular psychiatry 2012, 2012(17):290–314.

63. Lin MY, Sheng ZH: Regulation of mitochondrial transport in neurons. Exp Cell Res 2015, 334(1):35–44.

64. Shoffner J, Hyams L, Langley GN, Cossette S, Mylacraine L, Dale J, Ollis L, Kuoch S, Bennett K, Aliberti A et al: Fever plus mitochondrial disease could be risk factors for autistic regression. Journal of child neurology 2010, 25(4):429–434.

65. Poling JS, Frye RE, Shoffner J, Zimmerman AW: Developmental regression and mitochondrial dysfunction in a child with autism. Journal of child neurology 2006, 21(2):170–172.

66. Khemakhem AM, Frye RE, El-Ansary A, Al-Ayadhi L, Bacha AB: Novel biomarkers of metabolic dysfunction is autism spectrum disorder: potential for biological diagnostic markers. Metabolic brain disease 2017, 32(6):1983–1997.

67. Laszlo A, Horvath E, Eck E, Fekete M: Serum serotonin, lactate and pyruvate levels in infantile autistic children. Clinica chimica acta; international journal of clinical chemistry 1994, 229(1–2):205–207.

68. Essa MM, Guillemin GJ, Waly MI, Al-Sharbati MM, Al-Farsi YM, Hakkim FL, Ali A, Al-Shafaee MS: Increased markers of oxidative stress in autistic children of the Sultanate of Oman. Biological trace element research 2012, 147(1–3):25–27.

69. Gu F, Chauhan V, Kaur K, Brown WT, LaFauci G, Wegiel J, Chauhan A: Alterations in mitochondrial DNA copy number and the activities of electron transport chain complexes and pyruvate dehydrogenase in the frontal cortex from subjects with autism. Transl Psychiatry 2013, 3:e299.

70. Anitha A, Nakamura K, Thanseem I, Yamada K, Iwayama Y, Toyota T, Matsuzaki H, Miyachi T, Yamada S, Tsujii M et al: Brain region-specific altered expression and association of mitochondria-related genes in autism. Mol Autism 2012, 3(1):12.

71. Anitha A, Nakamura K, Thanseem I, Matsuzaki H, Miyachi T, Tsujii M, Iwata Y, Suzuki K, Sugiyama T, Mori N: Downregulation of the expression of mitochondrial electron transport complex genes in autism brains. Brain pathology (Zurich, Switzerland) 2013, 23(3):294–302.

72. Ginsberg MR, Rubin RA, Falcone T, Ting AH, Natowicz MR: Brain transcriptional and epigenetic associations with autism. PLoS One 2012, 7(9):e44736.

73. Scarpulla RC: Metabolic control of mitochondrial biogenesis through the PGC-1 family regulatory network. Biochimica et biophysica acta 2011, 1813(7):1269–1278.

74. Jornayvaz FR, Shulman GI: Regulation of mitochondrial biogenesis. Essays Biochem 2010, 47:69–84.

75. Ventura-Clapier R, Garnier A, Veksler V: Transcriptional control of mitochondrial biogenesis: the central role of PGC-1alpha. Cardiovasc Res 2008, 79(2):208–217.

76. Xu Y, Kabba JA, Ruan W, Wang Y, Zhao S, Song X, Zhang L, Li J, Pang T: The PGC-1alpha Activator ZLN005 Ameliorates Ischemia-Induced Neuronal Injury In Vitro and In Vivo. Cell Mol Neurobiol 2018, 38(4):929–939.

77. Zhang LN, Zhou HY, Fu YY, Li YY, Wu F, Gu M, Wu LY, Xia CM, Dong TC, Li JY et al: Novel small-molecule PGC-1alpha transcriptional regulator with beneficial effects on diabetic db/db mice. Diabetes 2013, 62(4):1297–1307.

78. Gotham K, Risi S, Pickles A, Lord C: The Autism Diagnostic Observation Schedule: revised algorithms for improved diagnostic validity. J Autism Dev Disord 2007, 37(4):613–627.

79. Lord C, Risi S, Lambrecht L, Cook EH, Jr., Leventhal BL, DiLavore PC, Pickles A, Rutter M: The autism diagnostic observation schedule-generic: a standard measure of social and communication deficits associated with the spectrum of autism. J Autism Dev Disord 2000, 30(3):205–223.

