## Supplemental Material for "Molecular Changes in Prader-Willi Syndrome Neurons Reveals Clues About Increased Autism Susceptibility"

Supplementary Material

### 1 Supplementary Tables

| **Cell Line** | **Diagnosis** | **Sex** | **ASD Status** |
| --- | --- | --- | --- |
| 38* | N/A | F | Unknown |
| 182* | N/A | F | Normal |
| 189* | N/A | F | Unknown |
| 238* | N/A | M | Normal |
| 195 | N/A | M | Normal |
| 312 | N/A | M | Normal |
| 152* | PW deletion | M | Normal |
| 192* | PW deletion | F | Normal |
| 225* | PW deletion | M | Normal |
| 258* | PW deletion | F | Normal |
| 148 | PW deletion | M | Normal |
| 307 | PW deletion | F | Normal |
| 156* | PW UPD | F | Normal |
| 162* | PW UPD | F | Normal |
| 171* | PW UPD | F | Normal |
| 249* | PW UPD | F | Normal |
| 270 | PW UPD | F | Normal |
| 191 | PW UPD | M | Normal |
| 235 | PW UPD | M | Normal |
| 197 | PW UPD | M | Normal |
| 94* | PW UPD | F | Possible ASD |
| 228* | PW UPD | M | Possible ASD |
| 250* | PW UPD | M | Possible ASD |
| 268* | PW UPD | F | Possible ASD |
| 277 | PW UPD | F | Possible ASD |
| 181 | PW UPD | M | Possible ASD |

**Supplementary Table 1**. Cell lines used for RNAseq and immunocytochemistry analysis. Asterisk (*) denotes subjects used in RNAseq experiment.

| **Gene** | **Function** | **Link to PWS** |
| --- | --- | --- |
| ***PEX10*** | Peroxisomes breakdown toxins and synthesize lipids | Mutations lead to Zellweger syndrome. Primary phenotypes of Zellweger syndrome are hypotonia and neurodevelopmental delay [80] |
| ***SNTB2*** | Plays a role in regulation of secretory granules and localization of membrane proteins | Dystrophins are associated with muscular dystrophy [81] |
| ***CTDP1*** | Dephosphorylates a subunit of RNApol making it available for transcription | Mutations are associated with neuropathy, facial dysmorphism, and cataracts [82] |
| ***AKT1S1*** | Subunit of mTORC, which regulates cell growth in response to nutrients and hormonal signaling | Altered mTORC signaling has been linked to a variety of neurological disorders including autism, epilepsy, and neurodegenerative disorders [83] |
| ***ATP7A*** | Supplies copper to copper requiring proteins in the secretory pathway | Associated with Menkes disease [84]. Primary phenotypes are hypotonia, hypothermia, failure to thrive, and seizures. |
| ***MYL5*** | Component of ATPase cellular motor protein myosin | Myosin has many neuronal functions, particularly at the synapse [85] |
| ***MIPOL1*** | Function is not well characterized | Link to hand and feet abnormalities |
| ***TMEM92*** | Transmembrane protein with unknown function | Unknown |
| ***DHRS1*** | Functions as an oxidoreductase | Associated with speech/ language impairment |

**Supplementary Table 2**. Core PWS transcripts outside of the 15q11.2-13.1 region that were significantly different versus control in all PWS subgroup.

| **Gene** | **Full Name** | **Function** |
| --- | --- | --- |
| ***ACOT9*** | Acyl-CoA Thioesterase 9 | Mitochondrial acyl-CoA thioesterase |
| ***AGPAT5*** | 1-Acylglycerol-3-Phosphate O-Acyltransferase 5 | Expressed in mitochondria |
| ***AIFM1*** | Apoptosis Inducing Factor Mitochondria Associated 1 | NADH oxidoreductase found in the mitochondrial intermembrane space |
| ***BNIP1*** | BCL2 Interacting Protein 1 | Involved in mitophagy |
| ***CLTC*** | Clathrin Heavy Chain | Expressed in mitochondria |
| ***CMC4*** | C-X9-C Motif Containing 4 | Expressed in mitochondria |
| ***COX5B*** | Cytochrome C Oxidase Subunit 5B | Component of the electron transport chain |
| ***COX7B*** | Cytochrome C Oxidase Subunit 7B | Component of the electron transport chain |
| ***COX7C*** | Cytochrome C Oxidase Subunit 7C | Component of the electron transport chain |
| ***CS*** | Citrate Synthase | TCA cycle enzyme |
| ***CYB5A*** | Cytochrome B5 Type A | Electron carrier expressed in mitochondria |
| ***EHHADH*** | Enoyl-CoA Hydratase And 3-Hydroxyacyl CoA Dehydrogenase | Involved in mitochondrial fatty acid oxidation |
| ***FDXR*** | Ferredoxin Reductase | Mitochondrial flavoprotein involved in the electron transport chain |
| ***GFM1*** | G Elongation Factor Mitochondrial 1 | Mitochondrial translation elongation factor |
| ***GHR*** | Growth Hormone Receptor | Expressed in mitochondria |
| ***GLUD1*** | Glutamate Dehydrogenase 1 | Mitochondrial matrix enzyme |
| ***GPX1*** | Glutathione Peroxidase 1 | Protects cells from oxidative stress |
| ***GSTP1*** | Glutathione S-Transferase Pi 1 | Protects cells from oxidative stress |
| ***HAP1*** | Huntingtin Associated Protein 1 | Expressed in mitochondria |
| ***MFN2*** | Mitofusin 2 | Involved in mitochondrial fusion and maintenance of the mitochondial matrix |
| ***MINOS1*** | Mitochondrial Contact Site And Cristae Organizing System Subunit 10 | Involved in the maintenance of mitochondrial architecture |
| ***MRPL4*** | Mitochondrial Ribosomal Protein L4 | Component of the mitochondrial ribosomal 39S subunit |
| ***MRPL41*** | Mitochondrial Ribosomal Protein L41 | Component of the mitochondrial ribosomal 39S subunit |
| ***MRPL47*** | Mitochondrial Ribosomal Protein L47 | Component of the mitochondrial ribosomal 39S subunit |
| ***MRPS33*** | Mitochondrial Ribosomal Protein S33 | Component of the mitochondrial ribosomal 28S subunit |
| ***MRPS5*** | Mitochondrial Ribosomal Protein S5 | Component of the mitochondrial ribosomal 28S subunit |
| ***MTIF2*** | Mitochondrial Translational Initiation Factor 2 | Mitochondrial translation intitation factor |
| ***NAXE*** | NAD(P)HX Epimerase | Involved in mitochondrial metabolic processes |
| ***NDUFA1*** | NADH:Ubiquinone Oxidoreductase Subunit A1 | Component of the electron transport chain |
| ***NDUFV2*** | NADH:Ubiquinone Oxidoreductase Core Subunit V2 | Component of the electron transport chain |
| ***NFS1*** | Cysteine Desulfurase, Mitochondrial | Expressed in mitochondria |
| ***NT5M*** | 5',3'-Nucleotidase, Mitochondrial | 5' nucleotidase localized to the mitochondrial matrix |
| ***OAT*** | Ornithine Aminotransferase | Mitochondrial enzyme ornithine aminotransferase |
| ***OXLD1*** | Oxidoreductase Like Domain Containing 1 | Expressed in mitochondria |
| ***PERP*** | P53 Apoptosis Effector Related To PMP22 | Expressed in mitochondria |
| ***POR*** | Cytochrome P450 Oxidoreductase | Enzyme involved in electron transport |
| ***PRKAR2B*** | Protein Kinase CAMP-Dependent Type II Regulatory Subunit Beta | Plays a role in regulating energy balance |
| ***RMND1*** | Required For Meiotic Nuclear Division 1 Homolog | Involved in mitochondrial translation |
| ***SDHC*** | Succinate Dehydrogenase Complex Subunit C | Component of the electron transport chain |
| ***SDHD*** | Succinate Dehydrogenase Complex Subunit D | Component of the electron transport chain |
| ***SLC25A36*** | Solute Carrier Family 25 Member 36 | Mitochondrial transport protein |
| ***SLC25A39*** | Solute Carrier Family 25 Member 39 | Mitochondrial transport protein |
| ***SLC25A5*** | Solute Carrier Family 25 Member 5 | Mitochondrial transport protein |
| ***SLC35F6*** | Solute Carrier Family 35 Member F6 | Involved in the maintenance of mitochondrial membrane potential |
| ***SUCLG2*** | Succinate-CoA Ligase GDP-Forming Subunit Beta | TCA cycle enzyme |
| ***SYNE2*** | Spectrin Repeat Containing Nuclear Envelope Protein 2 | Expressed in mitochondria |
| ***TIMM17A*** | Translocase Of Inner Mitochondrial Membrane 17A | Mitochondrial transport protein |
| ***TMEM160*** | Transmembrane Protein 160 | Expressed in mitochondria |
| ***TMTC1*** | Transmembrane O-Mannosyltransferase Targeting Cadherins 1 | Expressed in mitochondria |
| ***TP63*** | Tumor Protein P63 | Expressed in mitochondria |
| ***TUFM*** | Tu Translation Elongation Factor, Mitochondrial | Mitochondrial translation elongation factor |
| ***TXN*** | Thioredoxin | Involved in redox reactions |
| ***UQCR10*** | Ubiquinol-Cytochrome C Reductase, Complex III Subunit X | Component of the electron transport chain |
| ***UQCR11*** | Ubiquinol-Cytochrome C Reductase, Complex III Subunit XI | Component of the electron transport chain |
| ***UQCRC2*** | Ubiquinol-Cytochrome C Reductase Core Protein 2 | Component of the electron transport chain |
| ***UQCRFS1*** | Ubiquinol-Cytochrome C Reductase, Rieske Iron-Sulfur Polypeptide 1 | Component of the electron transport chain |
| ***UQCRH*** | Ubiquinol-Cytochrome C Reductase Hinge Protein | Component of the electron transport chain |
| ***USP30*** | Ubiquitin Specific Peptidase 30 | Involved in mitophagy |

**Supplementary Table 3.** List of genes identified in DAVID mitochondrial enrichment categories.

#### 1.1 Supplementary Figures

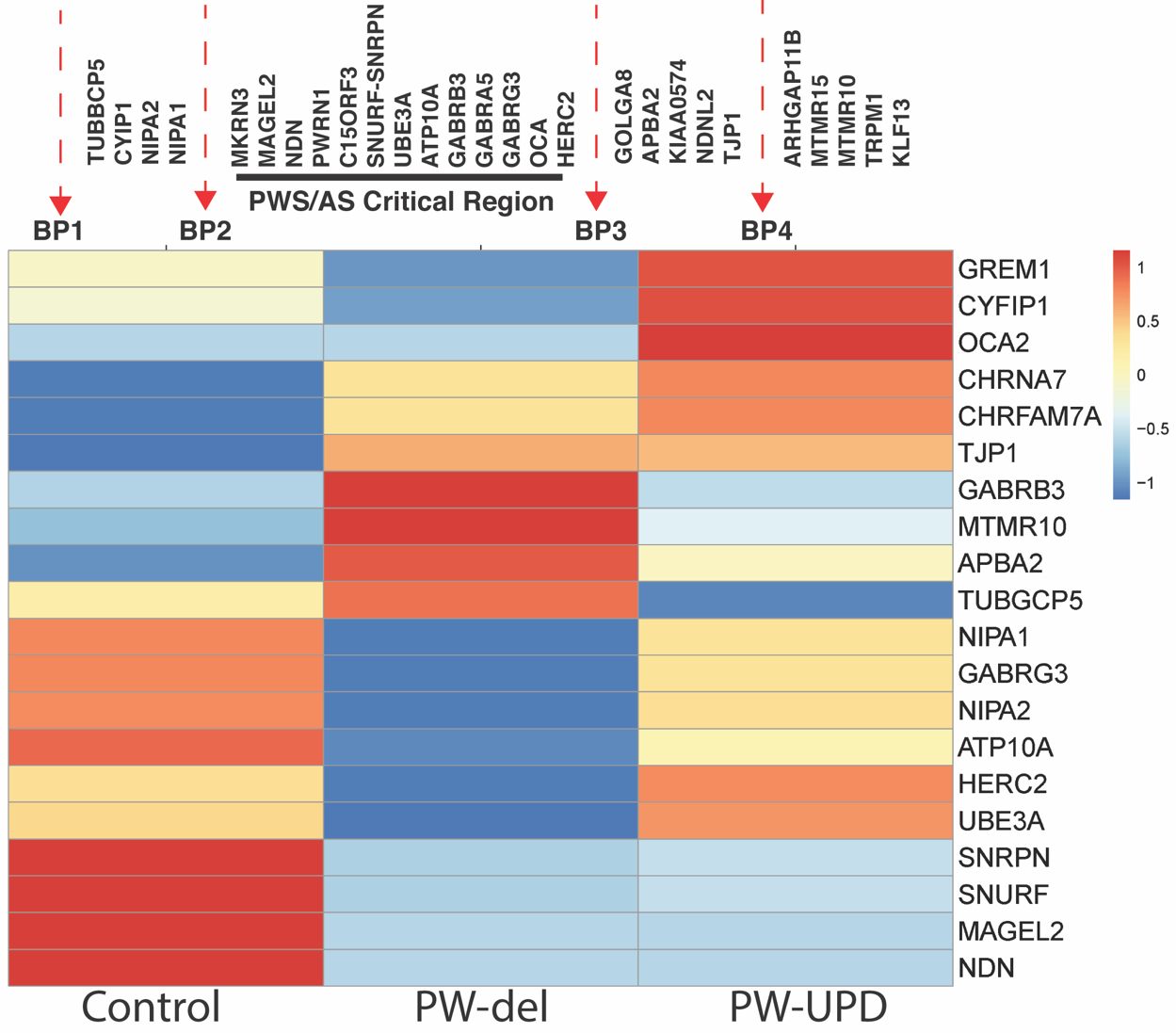

**Supplementary Figure 1. Genes in the 15q11.2-q13 critical region shows expected PWS imprinted expression.** Across the PWS/AS critical region, maternally imprinted genes such as *MAGEL2*, *SNRPN*, and *SNURF* showed decreased expression in both PW-del and PW-UPD neurons.

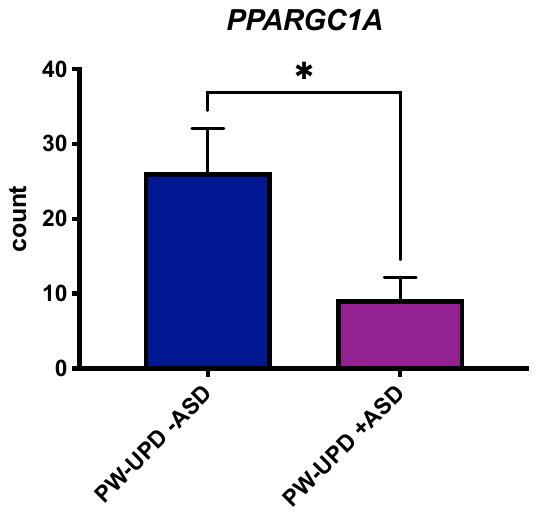

**Supplementary Figure 2. Expression of mitochondrial biogenesis factor, *PPARGC1A*, is significantly decreased in PW-UPD +ASD neurons.** Mean RNAseq count for each group is shown in the graph. Significance was determined by an unpaired *t*-test (p ≤ 0.05).
